# Transcriptome profiling of dopaminergic neurons derived from an ADHD induced pluripotent stem cell (iPSC) model

**DOI:** 10.1101/2024.09.22.614376

**Authors:** Atefeh Namipashaki, Aurina Arnatkeviciute, Shane D. Hellyer, Cameron J Nowell, Kevin Walsh, Jose M. Polo, Karen J Gregory, Mark A. Bellgrove, Ziarih Hawi

**Affiliations:** Turner Institute for Brain and Mental Health, School of Psychological Sciences, Monash University, VIC, Australia; Drug Discovery Biology, Monash Institute of Pharmaceutical Sciences, Monash University, Parkville, VIC, 3052, Australia; Department of Anatomy & Developmental Biology, Monash University, Melbourne VIC 3800, Australia. Development and Stem Cells Program, Monash Biomedicine Discovery Institute, Melbourne VIC 3800, Australia. Australian Regenerative Medicine Institute, Monash University, Melbourne VIC 3800, Australia; Adelaide Centre for Epigenetics and the South Australian Immunogenomics Cancer Institute, The University of Adelaide, SA, Australia

**Keywords:** ADHD, Dopaminergic differentiation, Stem cell modelling, Transcriptomics, Neurite outgrowth

## Abstract

The most recent ADHD***–***GWAS meta**–**analysis highlighted the potential role of 76 genes enriched among genes expressed in early brain development and associated with midbrain dopaminergic neurons. However, the precise functional importance of the GWAS-identified single nucleotide polymorphisms (SNPs) remain unknown. In contrast to GWAS, transcriptome analysis directly investigates gene products by assessing the transcribed RNA. This allows one to gain functional insights into gene expression paving the way for a better understanding of the molecular risk mechanisms of conditions, such as ADHD. In this study, we performed transcriptome profiling of highly homogeneous dopamine neurons developed from induced pluripotent cells (iPSCs) that were derived from an individual with ADHD and a neurotypical comparison individual. Comparative gene expression analysis between the examined lines revealed that the top differentially expressed genes (DEGs) were predominantly associated with nervous system functions related to neuronal development and dopaminergic regulation. Notably, 29 of the DEGs overlapped with those identified by ADHD***–***GWAS meta-analysis. These genes are overrepresented in biological processes including developmental growth regulation, axonogenesis, and nervous system development. In addition, gene set analysis revealed significant enrichment for meta categories such as ion channel activity, synaptic function and assembly, neuronal development and cell differentiation. Further, we observed significantly reduced projections in the ADHD dopamine neurons at the mid**–** differentiation stage (day 14 in vitro), providing preliminary support for the delayed neuronal maturation hypotheses of ADHD. This study underscores the potential of using iPSC**–**derived cell type**–**specific models that integrate genome and transcriptome analyses for biological discovery in ADHD.

## Introduction

Attention deficit hyperactivity disorder (ADHD) is the most prevalent neurodevelopmental condition, characterised by symptoms of inattention, impulsivity/hyperactivity or a combination of both (Fayyad et al., 2007). It affects 5% of school**–**age children globally and persists into adulthood in at least 60% of cases (Cherkasova et al., 2022). ADHD significantly impacts the life of affected individuals and their families with negative consequences on educational, occupational attainment, and reduced life expectancy (Barkley & Fischer, 2019). This multifaceted disorder is influenced by a complex interplay of genetic and environmental factors with estimated heritability ranging from 70% to 90% (Faraone et al., 2024; Levy, Hay, McStephen, Wood, & Waldman, 1997).

Psychostimulant medications that modulate catecholamine (dopamine and noradrenaline) neurotransmission have been the mainstay pharmacological treatment for ADHD since the 1950s. The effectiveness of these medications in reducing the core symptoms of ADHD led to the hypothesis that dysregulation of catecholamine signalling must be central to the aetiology of ADHD. This hypothesis has arguably been most extensively explored with respect to dopamine dysregulation across both human (e.g., molecular imaging of the dopamine transporter, DAT; candidate gene studies of the dopamine system) and animal studies (e.g., DAT1 knock**–**out in mice) (Giros, Jaber, Jones, Wightman, & Caron, 1996; Hawi et al., 2015; Krause, Dresel, Krause, Kung, & Tatsch, 2000). Inconsistencies across studies and methodological limitations have nevertheless prevented the identification of a robust signature of dopamine (or noradrenaline) dysfunction in ADHD. However, recent large GWAS meta-analysis has identified replicable associations between ADHD and 27 genetic loci encompassing 76 genes enriched in early brain development and associated with midbrain dopaminergic neurons (Demontis et al., 2023). Despite the success of GWAS in mapping risk loci for ADHD, the molecular insights derived from their findings remain limited. The challenges of interpreting GWAS findings can be attributed to the fact that neighbouring SNPs are often inherited together during meiotic recombination which makes it difficult to distinguish the causal SNPs underpinning the association (Cano-Gamez & Trynka, 2020). Further, the associated SNPs are often mapped between genes, adding another challenge in distinguishing the actual predisposing risk gene (Cano-Gamez & Trynka, 2020). In contrast, transcriptome analysis can accurately quantify gene expression levels and establish a global view of the whole transcriptome. Therefore, integrating genomic and transcriptomic findings in ADHD has the potential to extend our understanding of the molecular underpinnings of the disorder.

A key limitation within the field of psychiatric genetics, however, has been the lack of access to available brain tissue. For ADHD, recent studies have employed imputation techniques to predict gene expression of various brain regions based on genotype information, offering an indirect insight into ADHD**–**associated genes (Demontis et al., 2023; Liao et al., 2019). Others have examined peripheral blood and post**–**mortem brain tissue from individuals with ADHD and unaffected donors to identify differentially expressed genes and enriched pathways/networks relevant to the aetiology of the disorder (McCaffrey et al., 2020; Sudre et al., 2023). However, there are limitations to these approaches with blood cells lacking specificity for studying the transcriptomic profile of the condition, and post**–**mortem brain tissue composed of heterogeneous cell populations and being non-proliferative, making it unsuitable for culturing and expansion in the lab (Kampmann, 2020). Quantifying gene expression in neuronal cell types relevant to ADHD (i.e. dopaminergic neurons) is essential for understanding the mechanisms underlying the involvement of dopamine in the predisposition to ADHD. Thus, examining the transcriptome in dopaminergic neurons derived from ADHD and neurotypical comparison subjects represents a paradigm shift that could significantly advance our understanding of the aetiology of ADHD. Recently, a promising approach has been explored in psychiatric conditions utilizing patient**–**derived induced pluripotent stem cells (iPSCs) that can be differentiated into any cell type**–**specific context (e.g., dopaminergic neuron) (Matos, Ho, Schrode, & Brennand, 2020). This approach not only enables the integration of functional experiments to correlate altered transcriptomes with cellular function, but also offers the opportunity to study transcriptomic changes across different maturation timepoints.

In the current investigation, we performed transcriptome analysis using highly homogeneous dopamine neurons derived from ADHD and neurotypical comparison iPSCs. Differential gene expression analysis revealed significant under-expression of genes implicated in biological processes including regulating dopaminergic neurons, synapse formation, and cell adhesion. Our analysis was also extended to include transcriptomic changes with cellular phenotype by measuring neurite outgrowth during iPSC differentiation to dopaminergic neurons. Assessing neuronal maturity over the period of differentiation, contributes to further understanding the underlying biological mechanisms of ADHD.

## Methods and Materials

### Stem cell culturing

iPSCs derived from a male youth with ADHD and an unrelated male neurotypical comparison line (HDFn)(Namipashaki et al., 2023; Tong et al., 2019). The two lines were generated using CytoTune™**–**iPS 2.0 Sendai Reprogramming Kit (Invitrogen, Cat#A16517). Both lines were seeded on Biolaminin**–**521 (Biolamina, Cat#LN521) coated plates and maintained in StemFlex medium (Gibco, Cat#A3349401) for up to three passages. This allowed cells to adapt and reach a healthy state with a minimum of random differentiation.

### Differentiation of iPSCs into dopaminergic neurons

A lentiviral induction**–**based method was used to generate dopaminergic neurons (Powell et al., 2023). Briefly, three replicates for each of the iPSCs lines were differentiated via culturing with an inducible lentiviral system engineered to carry three transcription factors (*ASCL1***–***LMX1B***–** *NURR1*) and an antibiotic resistant (**–***PuroR)* gene. The cells were maintained on Biolaminin 111 LN (Biolamina, Cat#LN111) **–** coated plates in a DMEM/F12 (Gibco, Cat#10565018) based medium supplemented with Doxycycline (Sigma, #D9891) to induce the expression of the aforementioned genes. Puromycin (Sigma, #7255) was also added for selection of the cells that had successfully integrated the four genes within the genome. On DIV 14 (14 days *in vitro*), the media were changed to BrainPhys (Stem Cell Technologies, #05790), a neurophysiological culture medium known to improve neuronal function and supplemented with growth factors until maturated into dopamine neurons at day 35.

### Immunofluorescence detection of dopamine biomarkers

To verify that the developed neurons were dopaminergic, we examined them for the expression of dopamine neuron markers including tyrosine hydroxylase (TH; a rate**–**limiting enzyme of dopamine synthesis) and microtubule associated protein 2 (MAP2; a robust neuron**–**specific somato **–** dendritic marker). The culturing media on day 35 were aspirated, and the developed neurons were washed twice using PBS solution. These neurons were subsequently fixed using 4% para**–**formaldehyde fixative solution, (Electron Microscopy Sciences, #15170) for 15 minutes, followed by three PBS washes and 5 minutes of incubation in permeabilization buffer (0.5% Triton X**–**100 (Sigma, #T8787)). After three additional PBS washes, neurons were incubated for 30 minutes in blocking buffer (3% BSA, Sigma, #A7906) and then incubated with the primary antibodies (Recombinant Anti**–**Tyrosine Hydroxylase antibody, #ab75875; Anti**–**MAP2 antibody, #ab92434) for 2 hours at 37°C. This was followed by three PBS washes and subsequent incubation with the appropriate secondary antibodies (Abcam, #ab150081, #ab150174) for 1 hour at 37°C. A further three washing steps were performed with the last wash containing NucBlue live Cell Stain (Thermo, #R37605). The neurons were immediately imaged with the EVOS M5000 Imaging System (Thermo Fisher Scientific, Cat# AMF5000).

### Measuring cellular dopamine

Whole**–**cell dopamine ELISA analysis was performed to estimate dopamine concentration in the generated ADHD and comparison dopamine neurons. The neurons **–** each line with three biological replicates **–** were first imaged using the cell viability stain (Thermo# A15001) to count the area of viable cells. Subsequently, they were harvested using Accutase and spun at 1000 × g for 5 minutes at room temperature (RT). Media supernatants were completely aspirated, and cell pellets were flash**–**frozen in liquid nitrogen. The ELISA test was carried out using the Dopamine Research ELISA Kit from ALPCO (#17**–**DOPHU**–**E01**–**RES) as described by the manufacturers. Cells were lysed by sonication (Diagenode Bioruptor Plus), for 10 cycles (30sec on, followed by 30sec off for each cycle). Each sample was split into two technical triplicates. Absorbance at 450 nm was measured on a CLARIOstar Plus Microplate Reader (BMG LABTECH). Dopamine concentrations of the samples were calculated using Microplate Reader data analysis Software (MARS) and were normalized to µMols dopamine per relative fluorescent unit (RFU) of viable cells (µMols/RFU).

## Transcriptome analysis

### RNA Extraction, Library Preparation and sequencing

Following media removal, cells were washed twice with PBS and RNA was isolated using TRIzol reagent (Thermo, #15596026). The RNA samples were then purified using RNeasy Kit (Qiagen #74104). Quantification, quality control, library preparation and sequencing of RNA samples were performed at the Australian Genome Research Facility (AGRF). Following the initial quality control of the samples, 100 nanograms (ng) of RNA samples were used to prepare the library and perform the sequencing, as follows. Using the whole**–**transcriptome sequencing workflow, total RNA library was prepared via Illumina Ribo**–**Zero Plus comprising an initial step of rRNA depletion. Failure to remove rRNA before library preparation leads to the majority of sequencing reads being consumed by rRNA, thereby reducing the overall depth of sequence coverage. Following rRNA depletion, the remaining RNA molecules were fragmented into 150bp’s and underwent first and second cDNA synthesis utilizing reverse transcription. Completed cDNA samples were then indexed using adapter ligation and amplified by PCR to produce the libraries. Samples were then sequenced via the Illumina NovaSeq X Plus System with the depth of 50M reads. RNA sequencing was performed at the Australian Genomic Research Facility (AGRF). Primary sequence image analysis was performed using the NovaSeq Control Software (NCS) v1.2.0.28691 and Real Time Analysis (RTA) v4.6.7. RTA base calling was conducted using NovaSeq instrument computer. The Illumina DRAGEN BCL Convert 07.021.645.4.0.3 pipeline was used to generate the sequence data. The generated data met the AGRF quality control standards.

### Processing of RNA Sequences and Identification of differentially expressed genes

We applied an automated nf**–**core/rnaseq bioinformatic pipeline to process the read pairs obtained from the ADHD and neurotypical control RNA sequencing data (Harshil Patel, 2023). Briefly, Trim Galore v0.6.7 was applied to raw sequencing data to remove the adapter and poor**–**quality sequence reads (Felix Krueger, 2021). The bioinformatic tool STAR was used to align and annotate RNA sequences to the human reference genome GRCh38 with ENSEMBL version 109 gene annotations (Dobin et al., 2013). For RNA sequencing data, BAM file reads were converted to FASTQ format using Samtools v1.14 and merged with the unmapped gene reads (Patro, Duggal, Love, Irizarry, & Kingsford, 2017). Bioconductor package tximport was used to normalize reads against transcript length and generate gene-level counts (Soneson, Love, & Robinson, 2015). Differential gene expression analysis was then carried out using the Degust web application (Powell., 2019), utilising the limma**–**voom method (Law, Chen, Shi, & Smyth, 2014). Differentially expressed genes (DEGs) representing significant transcriptomic changes across the lines (FDR < 0.05) were included in volcano plots, collectively referred to as ADHD-DEGs. The top 10 DEGs with a high effect size of log2 fold change⩾2 was selected based on the highest significance according to FDR and used for further analysis.

#### Analysis of ADHD–DEG overlap with GWAS identified genes for psychiatric disorders

Psychiatric conditions often have shared genetic components. Therefore, we analysed whether ADHD-associated DEGs overlapped with those implicated in major psychiatric conditions. To perform this, we employed a tiered approach. First, we focused on identifying shared genes with substantial effect sizes (FDR ⩽ 0.05 and log2 FC ⩾ 2). While stringent thresholds provide high confidence, they may exclude biologically meaningful genes with moderate to subtle changes. Therefore, to conduct a more comprehensive analysis, we also searched for genes with more subtle but still significant changes (FDR ⩽0.05), which may reveal subtle regulatory effects shared across psychiatric conditions. To systematically identify genes involved in major psychiatric disorders we used data derived from GWASs [ADHD (Demontis et al., 2023), schizophrenia (SCZ) (Trubetskoy et al., 2022), major depression (MDD) (Als et al., 2023), bipolar disorder (BD) (Mullins et al., 2021), and autism spectrum disorder (ASD) (Grove et al., 2019)]. SNPs were mapped to corresponding genes, based on their position on the DNA using MAGMA software package (de Leeuw, Mooij, Heskes, & Posthuma, 2015). The corresponding annotation file MAGMAdefault.genes.annot and reference panel (1000 Genomes European ancestry) for linkage disequilibrium (LD) calculation was downloaded from https://github.com/thewonlab/H-MAGMA. Gene analyses in MAGMA are based on a multiple linear principal component’s regression model where the gene *p**–***values are computed using an F**–**test. As a result, genes with a more significant involvement in a GWAS were assigned lower *p**–***values. To select a list of genes for each GWAS, we controlled the family**–** wise error at 0.05 using Bonferroni correction (by setting a gene *p**–***value threshold equal to 0.05 divided by the total number of identified genes in MAGMA analysis). The number of selected genes for psychiatric disorder GWASs included 95 genes for ADHD, 336 genes for BD, 465 genes for MDD, 922 genes for SCZ, and 14 genes for ASD. The degree of correspondence between ADHD**–**DEGs and GWAS**–**implicated genes for each disorder was assessed by evaluating the number of ADHD**–**DEGs present in the GWAS**–**identified list. The statistical significance of the overlap was evaluated through permutation testing by comparing the number of overlapping genes in the empirical data to an ensemble of 100 000 gene lists where a set of GWAS**–**identified genes (of the same size) was selected at random. The list of genes that significantly overlapped with ADHD**–**GWAS was subsequently submitted to assess their overrepresentation in the biological processes using www.geneontology.org (Ashburner et al., 2000) and to the STRING database for investigation of protein-protein interactions. The p-value in STRING’s PPI network is calculated by comparing the actual number of observed interactions to the expected number of interactions under a null hypothesis, assuming that proteins are connected at random (Szklarczyk et al., 2023).

### Enrichment analysis using gene score resampling (GSR)

GSR analysis examines whether functionally related groups of genes [e.g., annotated using Gene Ontology (GO)] contain the highest**–**scoring genes. Here, we used GRS to investigate what GO categories contained the most significantly differentially expressed genes. Each gene from the RNA**–**seq analysis was assigned a **–**log10(p_FDR_) score quantifying its involvement in differential expression. As a result, genes with a more significant differential expression were assigned higher values. We also performed a separate enrichment analysis for the under and over**–**expressed genes using all gene scores quantified as the: i) sign of the log**–**fold change times **–** log10(p_FDR_) to evaluate the over**–**expressed genes; and ii) inverse sign of the log**–**fold change times **–** log10(p_FDR_) to evaluate the under**–**expressed genes. Functional gene group analyses were performed using version 3.2 of ErmineJ software (Gillis, Mistry, & Pavlidis, 2010). Gene ontology annotations were obtained from GEMMA https://gemma.msl.ubc.ca/arrays/showArrayDesign.html?id=735 as Generic\_human\_ ncbiId s\_noParents.an.txt.gz on March 4, 2024 (last updated 2024**–**03**–**01) (Zoubarev et al., 2012). Gene Ontology terms and definitions were automatically downloaded by ErmineJ on March 4, 2024 as go.obo (data version 2024**–**01**–**17) and can be downloaded from http://release.geneontology.org/2024-01-17/ontology/index.html. We performed gene score resampling (GSR) analysis on the **-** log10(p_FDR_) scores testing the biological processes, molecular function, and cellular component GO categories with 5**–**100 genes available using the mean **–** log10(p_FDR_) score across genes to summarize the GO category and applying full resampling with 10^6^ iterations. These analyses were also repeated for under and over**–** expressed genes using signed scores as described above. The resulting *p***–**values were corrected across 9177 GO categories, controlling the false discovery rate (FDR) at 0.05 using the method of Benjamini and Hochberg (Benjamini, Drai, Elmer, Kafkafi, & Golani, 2001). The top 50 GO categories with the most significant FDRs were subsequently grouped into meta**–**categories based on their functional similarities.

## Neurite outgrowth measurement

For each sample, neurite outgrowth of the developing dopamine neurons was measured over the time progression of differentiation on days 7, 14 and 42. The culturing media were half changed three times with BrainPhys without phenol Red with the last change containing 2X concentration of cell viability stain (neurite outgrowth staining kit, Thermo# A15001). Following incubation for 15 minutes at 37^°^C, the developing neurons were imaged using Leica DMi8 fluorescence microscopy. The images were loaded into ilastik for pixel classification which uses an interactive machine learning bioimage algorithm for segmentation of cell bodies and neurites (Berg et al., 2019). An algorithm was generated for each timepoint and applied to analyze all images from that specific day. The probability map of the segmented images was processed using the Fiji image processing package to calculate the areas of neurites and cell bodies (Schindelin et al., 2012). The neurite’s outgrowth of ADHD cell lines was compared to those of the neurotypical comparison using the calculated total area of neurites relative to cell bodies for each time point. Statistical differences between the groups were evaluated using Student’s *t*-tests, with Bonferroni correction applied to account for multiple comparisons.

## Results

### Differentiation of iPSCs into dopaminergic neurons

The direct dopaminergic differentiation method produced highly homogeneous populations of dopaminergic neurons within 35 days. This method benefits from the integration of the exogenous transcription factors including *ASCL1***–***LMX1B***–***NURR1* (known to function in dopamine neurogenesis) together with the antibiotic resistance gene (*PuroR*), to successfully increase the homogeneity of the resulting neurons (Caiazzo et al., 2011; Powell et al., 2023). We also utilised the dopaminergic-specific extracellular matrix Laminin 111LN instead of the previously applied mouse sarcoma extracted matrix which was found to improve the homogeneity of dopamine neurons derived from iPSCs (Nolbrant, Heuer, Parmar, & Kirkeby, 2017). Consequently, counting the percentage of TH-positive neurons relative to the total number of cells positive for DAPI nuclei and MAP2 neuronal marker in the stained fields of views revealed highly homogeneous dopaminergic neurons with >99% purity by DIV35 (Figure 1A). Further, transcriptomic analysis confirmed the expression of ventral midbrain floor plate identity and specification markers *NR4A2*, *LMX1A, OTX2, ASCL1, MSX1* and *EN2,* in addition to dopaminergic maturation markers *TH, DDC, SLC6A3, SLC18A2, SYN1* and *DRD2* (Figure 1B). Measurement of the whole **–** cell dopamine in the resultant neurons on DIV35 showed a robust biosynthesis with a mean of 18.2 µMols/RFU dopamine across the cell lines. This provided further confirmation for the specificity and maturity of the differentiated midbrain dopaminergic neurons (Figure 1C).

**Figure 1.**
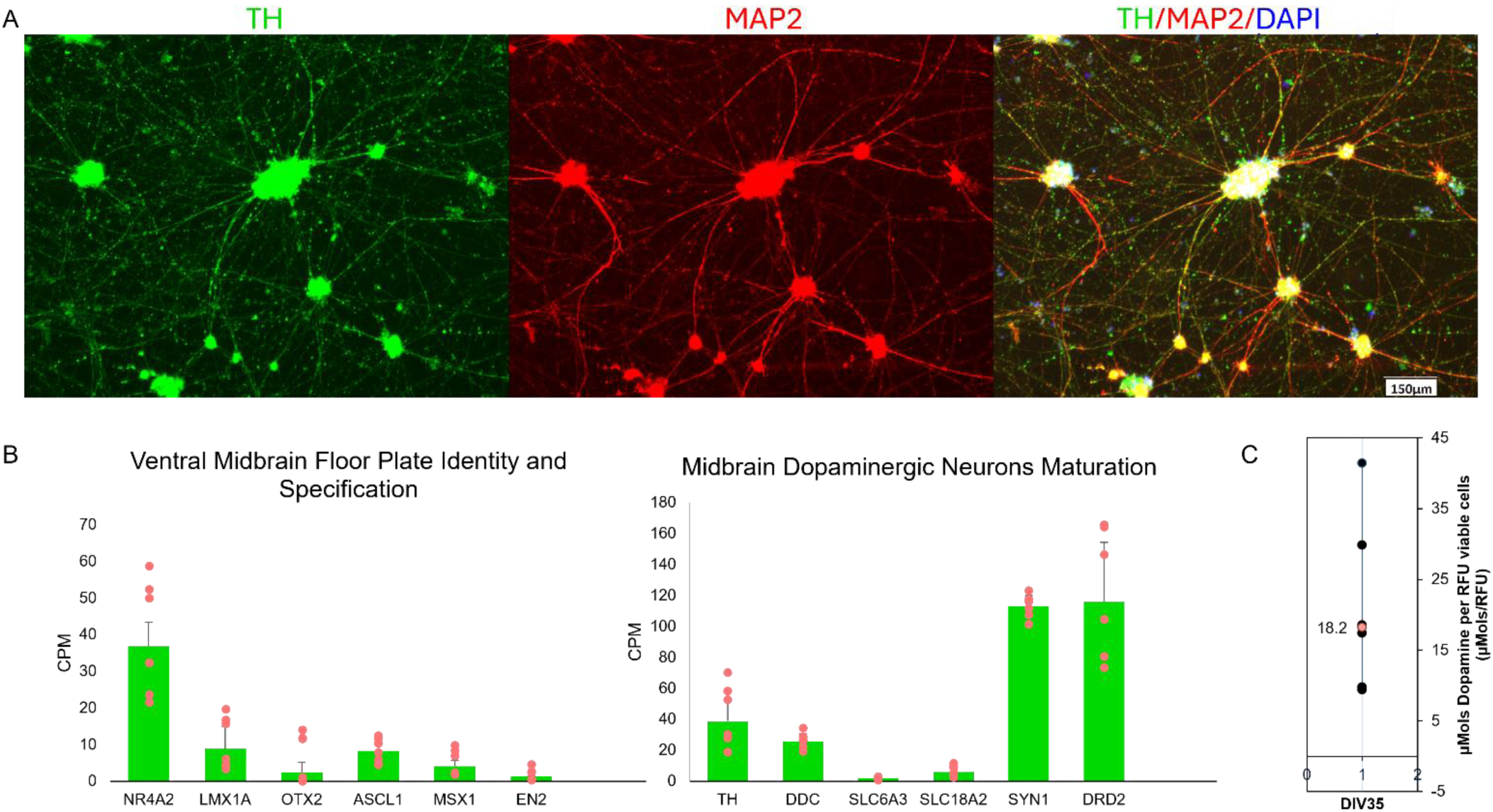
Confirmation of specificity and maturity of dopaminergic neurons. **(A)** Immunocytochemical staining of dopaminergic neurons showing expression of *TH* and *MAP2* and DAPI nuclear markers. Scale bar: 150µm (**B**) Expression of *NR4A2, LMX1A, OTX2, ASCL1, MSX1* and *EN2* confirms the dopamine neuron’s identity and specification, while the expression of *TH, DDC, SLC6A3, SLC18A2, SYN1* and *DRD2* demonstrates their maturity (**C**) ELISA test confirming dopamine biosynthesis and exhibiting a mean expression level of 18.2 µMols per RFU viable cells (µMols/RFU) at DIV35. RFU: Relative Fluorescent Unit.

### DEGs in ADHD dopaminergic neurons

Whole transcriptome RNA sequencing analysis showed that the total number of detected genes was 23,046, which is 99.98% identical to those reported using large scale RNA**–**Seq studies, indicating the reliability of the generated RNA**–**Seq data (Garcia-Ortega & Martinez, 2015). Quantifying gene expression differences between ADHD-derived dopamine neurons and those developed from a neurotypical control showed 891 genes with evidence of differential expression at false discovery rate (FDR) of ⩽ 0.05 and absolute log_2_ fold change (FC) of ⩾ 2 (Figure 2 A). Of these, the top 10 protein coding DEGs were found to have a highly significant absolute log_2_ FC ranging from 2.02 to 8.93 with FDR value of 1.06 x 10^-9^ to 2.15 x 10^-14^. Interestingly, these genes were either previously linked to major psychiatric disorders and neurological diseases (Parkinson’s and Alzheimer’s diseases) or found to be implicated in neurobiological functional categories central to the development of these conditions (Figure 2B, Table 1). These include dopaminergic regulation (*INPP5F, CBLN1, DPP6, DPP10* and *EPHA5*) (Cao, Park, Wu, & De Camilli, 2020; Cooper, Kobayashi, & Zhou, 2009; Iida et al., 2024; Jerng, Lauver, & Pfaffinger, 2007; Kaulin et al., 2009; Maffie & Rudy, 2008; Mazzocchi et al., 1999), neuronal development (*RBFOX1, PCDHA2, EPHA5, PTPRT, LARGE1* and *RTL1*) (Cooper et al., 2009; Kitazawa, Sutani, Kaneko-Ishino, & Ishino, 2021; Lee, 2015; Longman et al., 2003; Shao et al., 2019; W. W. Zhao, 2013), synapse regulation (*INPP5F, DPP6* and *PTPRT*) (Lee, 2015; Malloy, Ahern, Lin, & Hoffman, 2022; Nakatsu et al., 2015), animal behavioural alteration (*PTPRT, RBFOX1* and *RTL1*) (Kitazawa et al., 2021; Lee, 2015; O’Leary et al., 2022), and neuronal excitability (*RTL1*) (Chou et al., 2022). Dysregulation of these processes has been consistently reported to associate with the development of ADHD (Faraone & Larsson, 2019; Hawi et al., 2015). Further, RNA-seq data revealed 49 differentially expressed cadherin genes with FDR⩽0.05, eight of which (PCDHA2, PCDHA3, PCDHB5, PCDH20, CDH6, PCDHB17P, PCDHB18P) had a prominent absolute log2 fold change of ⩾2 (Supplementary table 1). Several genetic association studies have implicated cadherin superfamily genes as risk factors to the development of major psychiatric conditions including ADHD (Hawi et al., 2018).

**Figure 2.**
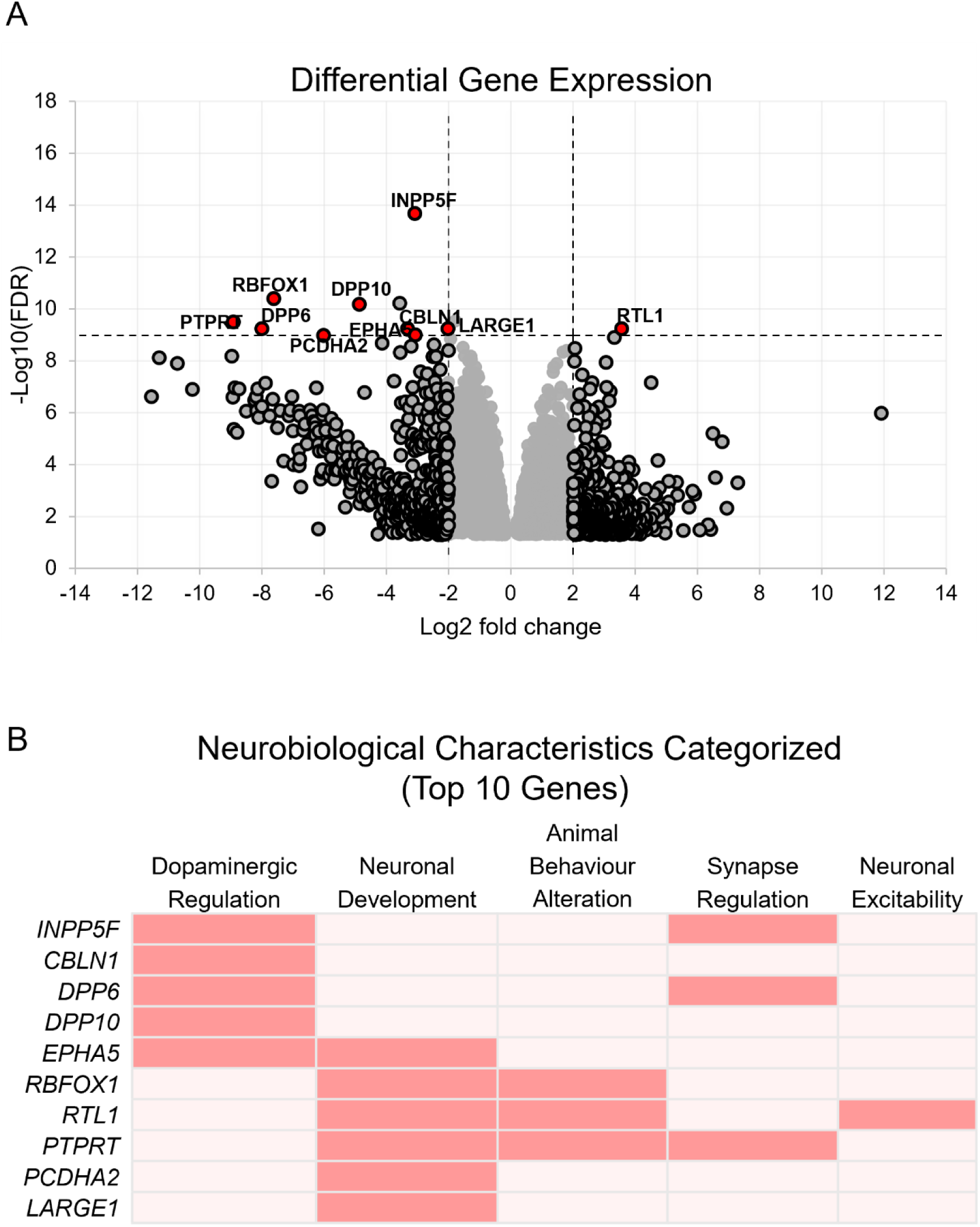
Differentially expressed genes in ADHD. (**A**) Volcano plot showing DEGs with FDR ⩽0.05. The horizontal line represents the cut off for the top10 protein-coding genes which are highlighted in red. Vertical lines indicate log2 fold change greater than ±2 (**B**) Heatmap table showing the meta categories of the neurobiological characteristics of these genes.

**Table 1.**
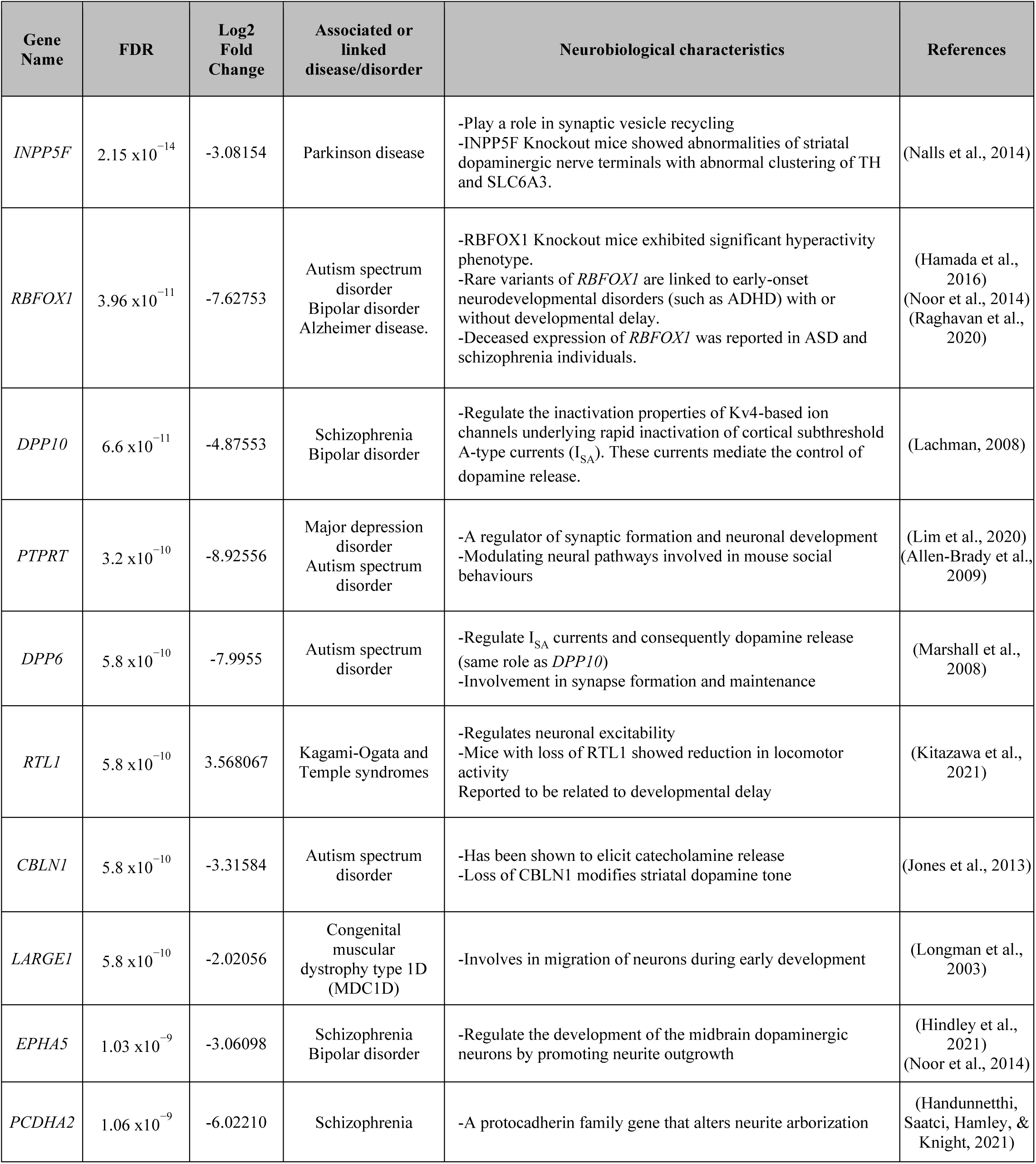
Neurobiological characteristics of the top 10 protein coding DEGs, their association with neurological/psychiatric conditions, FDR and Log2 fold change values.

#### ADHD–DEGs overlap with GWAS identified risk genes of major psychiatric disorders

As psychiatric conditions significantly overlap at symptom level and share significant genetic components, we used GWAS summary statistics to examine the extent to which ADHD**–**DEGs overlapped with GWAS**–**risk genes for the major psychiatric disorders. Two lists (Supplementary table 2 and 3) of the ADHD**–**DEGs, one with FDR⩽0.05 and the other with a more stringent list of FDR ⩽0.05 and log_2_ FC⩾2 were used to examine the potential overlap with ADHD, ASD, BD, MDD and SCZ genes identified using MAGMA software package (see Methods). More stringently derived ADHD-DEGs did not show a significant overlap with any of the GWAS-implicated genes reported for the five major psychiatric disorders after Bonferroni correction (p⩽0.01). However, DEGs with FDR of ⩽0.05 showed significant overlap with ADHD, MDD and SCZ, indicating shared genetic links between these conditions (Figure 3A). More specifically, 29 differentially expressed genes significantly overlapped with ADHD**–**GWAS associated genes (Supplementary table 4). Enrichment analysis of these genes revealed their overrepresentation in three biological processes, all related to neurodevelopment including regulation of axonogenesis (GO:0050770, FDR = 4.66 x 10^-2^), regulation of developmental growth (GO:0048638, FDR = 3.19 x 10^-2^), and nervous system development (GO:0007399, FDR = 3.70 x 10^-2^) (Figure 3B). Additionally, protein**–**protein interaction network analysis of these 29 genes showed a significantly higher number of interactions than expected under the null hypothesis, with a *p**–***value of 4.38 x 10^-5^, indicating non**–**random connections and significant number of edges (Figure 3C). The symptom overlap among psychiatric disorders was supported by the overlap of our ADHD**–**DEGs with GWAS-identified genes for ADHD, MDD, SCZ indicating a shared biological similarity among these conditions.

**Figure 3.**
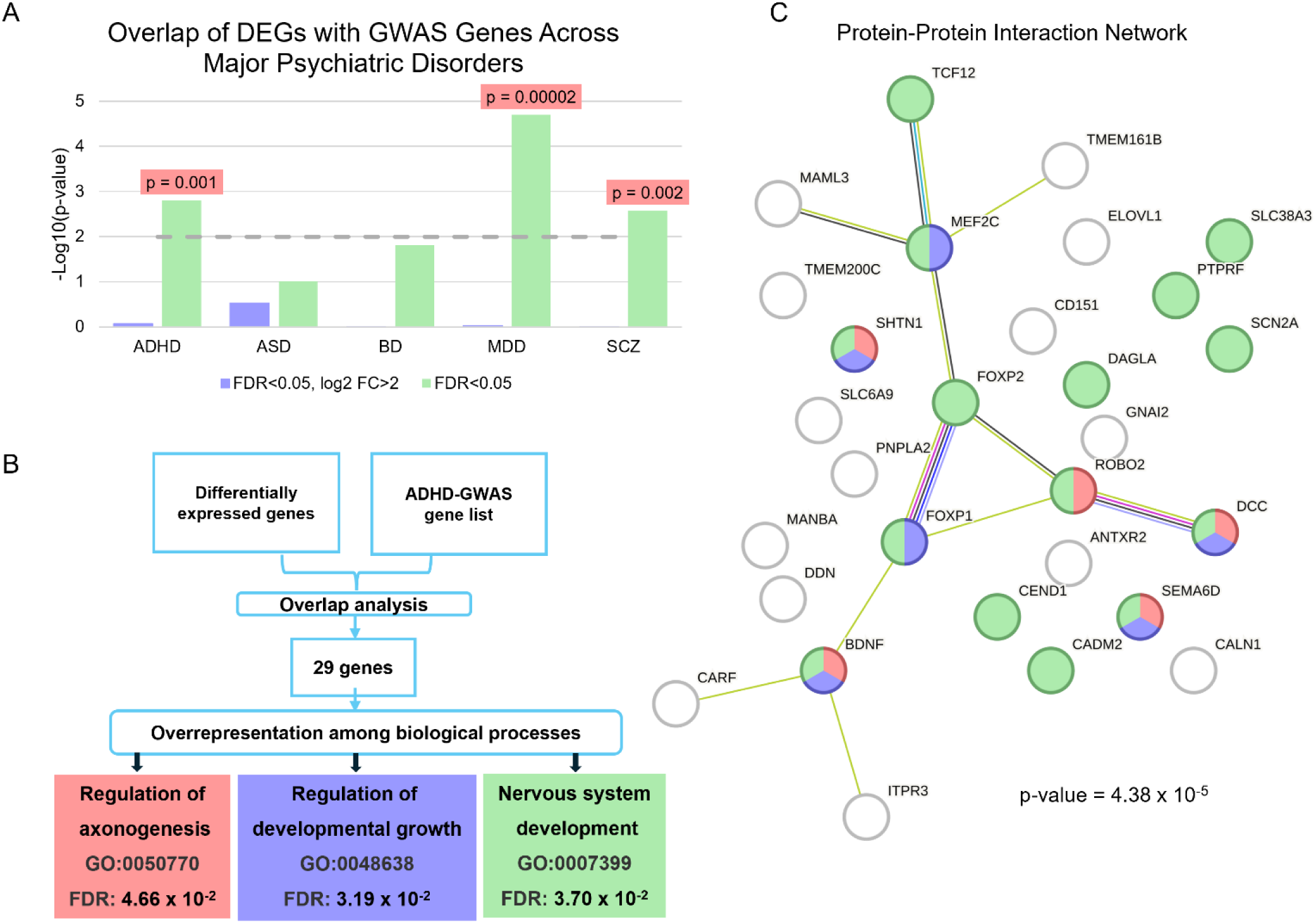
ADHD **–**DEGs overlap with GWAS findings for the 5 major psychiatric disorders. (**A**) A diagram showing the statistical significance of the overlaps (Bonferroni corrected p ⩽0.01) with ADHD, ASD, BIP, MDD and SCZ GWAS. The dotted line shows the cut**–**off for significance. Significant *p* values are highlighted in red. (**B**) A workflow analysis showing significant overrepresentation of ADHD **–** DEGs (n = 29) in biological processes including axonogenesis, neuronal growth and development. (**C**) STRING database show protein**–**protein interaction network of the 29 DEGs, displaying a significant number of interactions (p= 4.38 x 10^-5^). Colours match the represented Go term in the panel B.

### GO term enrichment analysis

Gene set enrichment analyses using GSR indicated that the top 50 gene ontology annotations with FDR ⩽ 0.01 mainly involved GO categories linked to neural processes (Supplementary table 5). Further grouping of these terms showed significant enrichment for meta categories including ion channel activity and signalling, synaptic function and assembly, neuronal development and differentiation, glutamate receptor activity, cellular signalling pathways, membrane and structural components, cell adhesion and migration and neurotransmitter activity (Figure 4A). While analysing the enrichment of all DEGs together can provide a broad understanding of biological processes, pathways and functions that are affected by the condition, it cannot show if certain processes/pathways are being activated or suppressed. Therefore, we performed separate enrichment analyses for the downregulated and upregulated genes. The enrichment of under**–**expressed genes resulted in similar meta**–**categories to those of total DEGs, except of those that are related to chromatin and gene regulation (Figure 4A and Supplementary table 6). However, evaluation of overexpressed genes showed only 6 GO**–**terms with a neural theme (Figure 4B and Supplementary table 7) including dopamine catabolic process (GO:0042420, *p* = 3 x 10⁻^3^), catecholamine catabolic process (GO:0042424, *p* = 3 x 10⁻^3^), cadherin binding involved in cell**–**cell adhesion (GO:0098641, *p* = 3 x 10⁻^3^), positive regulation of receptor binding (GO:1900122, *p* = 3 x 10⁻^3^), positive regulation of cell**–**matrix adhesion (GO:0001954, *p* = 3 x 10⁻^3^) and extracellular matrix binding (GO:0050840, *p* = 1.53 x 10⁻^9^). The results indicate that the downregulated genes are broadly involved in neural processes, similar to the overall DEGs, suggesting a general suppression of neural-related functions. However, the upregulated genes were more selectively enriched, primarily highlighting specific pathways such as dopamine catabolism, cell adhesion, and extracellular matrix interactions.

**Figure 4.**
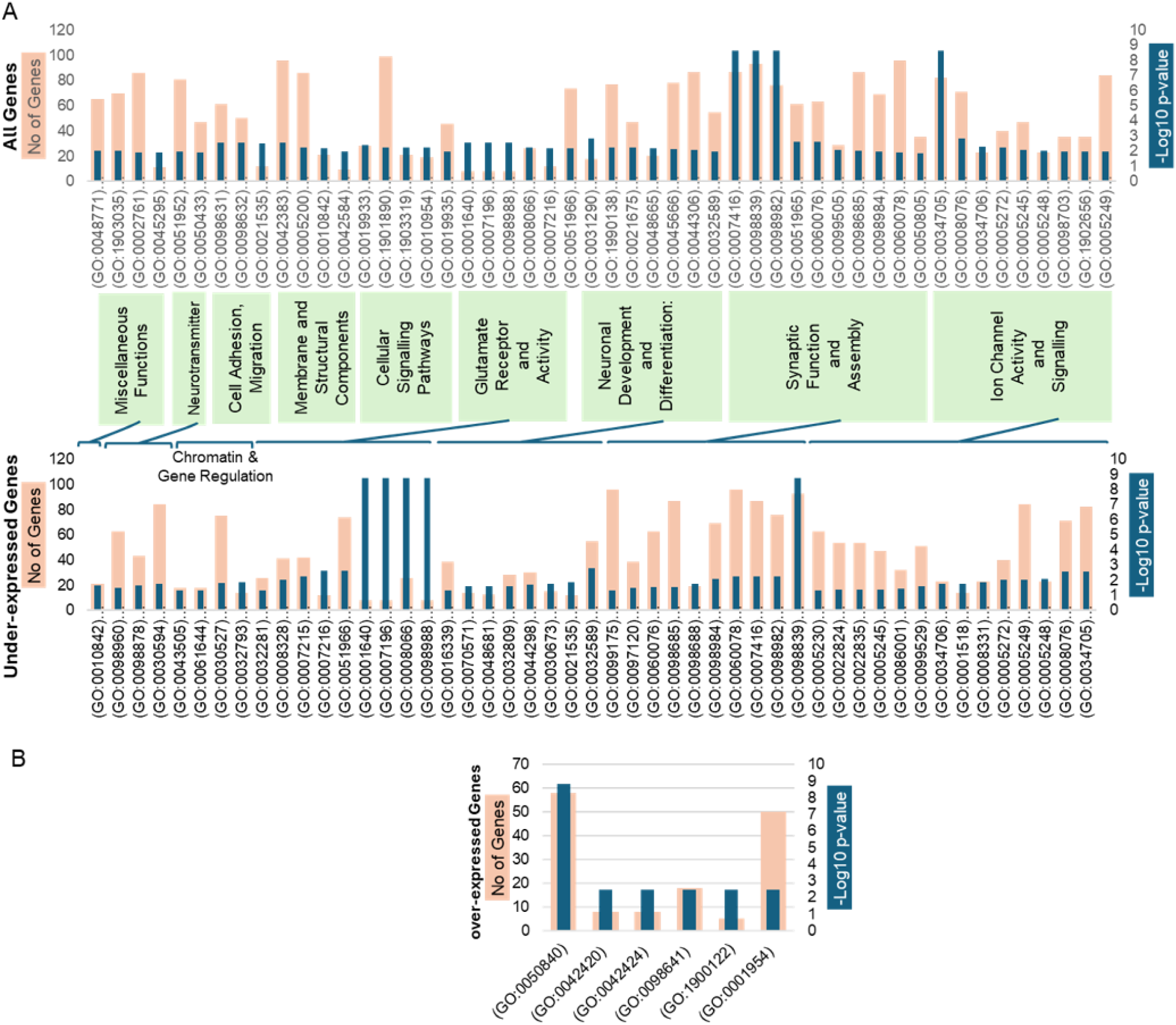
Gene enrichment analysis showing the top 50 enriched GO categories. (**A**) Enrichment result for the top 50 GO categories using total genes and under**–**expressed genes shows similar pattern for meta categories except of the chromatin and gene regulation. (**B**) Enrichment result for the top 50 GO categories using over**–**expressed genes shows only 6 significant GO term with the neural theme including dopamine catabolic process (GO:0042420), catecholamine catabolic process (GO:0042424), cadherin binding involved in cell**–**cell adhesion (GO:0098641), positive regulation of receptor binding (GO:1900122), positive regulation of cell**–**matrix adhesion (GO:0001954) and extracellular matrix binding (GO:0050840).

### Neurite outgrowth analysis of dopaminergic neurons

The consistent finding of neuronal development and differentiation in our association/ enrichment analyses of ADHD–DEGs underscores the brain maturational delay hypothesis of ADHD (Shaw et al., 2007). To explore this, we conducted neurite outgrowth analysis during the progression of dopamine neuron differentiation at three timepoints: 1) days in vitro (DIV)7: when the developing cells are at neuron progenitor stage and just beginning to elongate the neurites from the cell bodies (still in the lag phase of neurite outgrowth); 2) DIV14: when the cells are at mid–differentiation stage and are extending their neurites for maturation/completion of differentiation (in the exponential phase of neurite outgrowth); and 3) DIV42: the cells are at a maturational stage that has already passed the plateau phase of neurite outgrowth (due to confined cultureware area) and showing extensive branching and lengthy neurites. This assay allowed us to assess a key characteristic of neuronal development which is the extension of neural projections during differentiation/maturation. Fluorescent images of the cells stained for both cell bodies and neurites were obtained to create their segmentation map, enabling the calculation of the ratio of neurites to cell bodies at each timepoint. At DIV7, there were no significant differences in cell branching of the ADHD and neurotypical comparison (Figure 5A). However, we observed significantly fewer neuronal projections in the ADHD lines compared to the neurotypical comparison lines at the mid–differentiation stage on DIV14 (p = 0.008) (Figure 5B). By DIV42 no significant difference was observed in the neurite to cell body ratio (Figure 5C).

**Figure 5.**
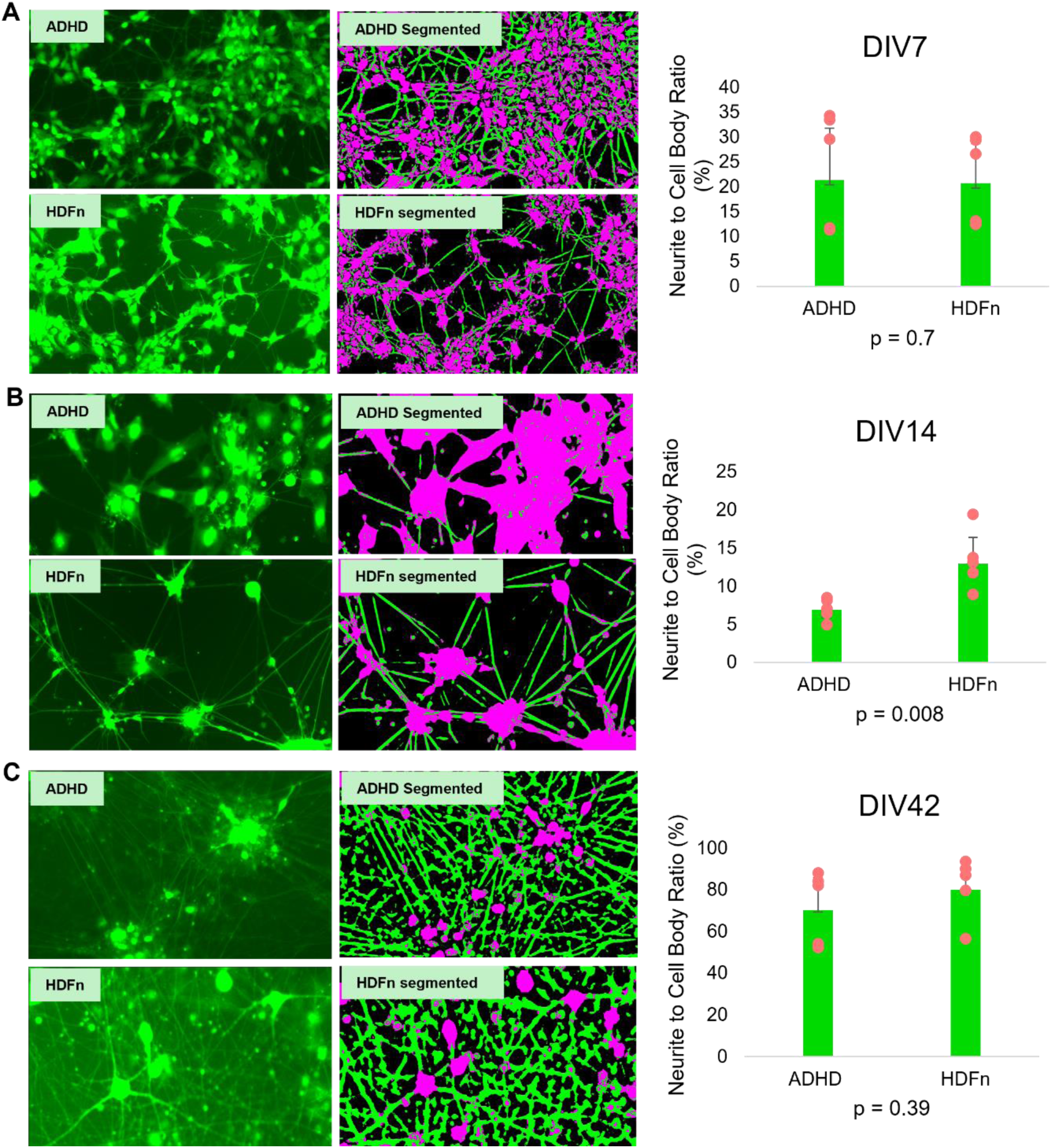
Neurite outgrowth assay. (**A**) Neuronal branching analysis showing no statistical differences at DIV 7 of iPSC differentiation to dopaminergic neurons. (**B**) Midway (DIV14) differentiation showing significant difference (p = 0.008) in neuronal projections. The ADHD line shows fewer neurite outgrowth compared to the comparison HDFn line. (**C**) On DIV42, the neurons of ADHD and comparison lines show extensive branching with no statistical differences between the two lines. The pink colour represents the neuronal cell bodies, while the green indicates the neurites in the segmented images.

Neurite outgrowth, similar to the growth curve observed in cultured cells, follows a multistage process with distinct phases. In ADHD neurons, we observed a delayed progression along this curve. No significant differences were found during the early lag phase (DIV7), but ADHD neurons entered the logarithmic growth phase later than neurotypical comparison (DIV14), resulting in fewer neurite projections at that stage, indicating a slower maturation process. By the maturation stage (DIV42), when neurite outgrowth had plateaued and filled the available space in the culture well, these differences appeared to resolve. However, further delays may still be present but are difficult to detect once the growth curve reaches the plateau phase.

## Discussion

This study presents transcriptome profiling of highly homogeneous ADHD dopaminergic neurons derived from iPSCs. The findings offer valuable insights into the molecular mechanisms underlying ADHD and underscore the importance of using pure, maturated iPSC- derived dopaminergic neurons for mechanistic studies of the condition. The high level of dopamine neuron homogeneity (99%) minimized variability, ensuring that the subsequent analyses accurately reflected dopaminergic neuron-specific characteristics (Muzes & Sipos, 2016). Systematic analysis of differentially expressed genes (DEGs) revealed that both the top-ranking and overall genes were strongly associated with and enriched in processes linked to the aetiology of ADHD. Additionally, in vitro phenotypic analysis demonstrated that ADHD neurons exhibited significantly fewer neuronal projections at DIV14, further supporting the role of these molecular alterations in the condition.

All of the top ten protein-coding DEGs are associated with biological functions relevant to the aetiology of ADHD. For example, knockout of the most significant DEG, *INPP5F*, in animal models has been shown to cause abnormal clustering of tyrosine hydroxylase (TH) and dopamine transporter (DAT) in striatal dopaminergic nerve terminals, both of which play crucial roles in regulating dopamine transmission at the synapse (Cao et al., 2020). *INPP5F* functionally overlaps with *SYNJ1* (synaptojanin 1), a gene involved in synaptic transmission and membrane trafficking, with mutations in *SYNJ1* leading to Parkinson’s disease, a condition characterized by low dopamine levels (Olgiati et al., 2014). Additionally, *INPP5F* has been identified as a GWAS risk gene for Parkinson’s disease (Blauwendraat, Nalls, & Singleton, 2020). The three-fold reduction in *INPP5F* expression observed in our ADHD samples may contribute to altered regulation of dopamine transmission. However, further empirical evidence is needed to confirm this hypothesis.

Gene set enrichment analyses have also revealed strong enrichment with biological functions relevant to the aetiology of ADHD. This includes the most significantly enriched GO term, ’synapse assembly’ (GO:0007416, p = 2.29 x 10⁻⁹), which comprises 87 ADHD DEGs. Disruption of synapses was recognised as the most commonly identified effect of ADHD - associated DNA variations (Dark, Homman-Ludiye, & Bryson-Richardson, 2018). Notably, our RNA-seq data showed a significant 8.9 log fold reduction in *PTPRT* expression, a brain-specific gene regulating synapse formation via cadherins (Arikkath & Reichardt, 2008). This, combined with alterations in 49 differentially expressed cadherin genes in our RNA seq data, suggests disrupted synapse formation potentially contributing to ADHD risk. Another enriched GO term category involves ion channel activity and signalling, which is strongly linked to ADHD. Both potassium and calcium channel genes have been reported to regulate dopamine transmission and associated with ADHD and other psychiatric conditions (Cross-Disorder Group of the Psychiatric Genomics, 2013; Fulton et al., 2011).

Our findings also highlight an enrichment of glutamatergic and GABAergic GO terms, aligning with the excitation-inhibition imbalance proposed in ADHD (Selten, van Bokhoven, & Nadif Kasri, 2018). Copy number variations (CNVs) in glutamate receptor genes (*GRM7, GRM8, GRM1*) have been reported in ADHD, with CNVs linked to cognitive and clinical characteristics (Akutagava-Martins et al., 2014; Elia et al., 2011). Additionally, studies have associated GABAergic genes, such as *GAD1*, with increased susceptibility to the hyperactivity and impulsivity symptoms of ADHD (Ferranti, Luessen, & Niswender, 2024). Animal models also support the role of disrupted glutamatergic transmission in ADHD, particularly in the prefrontal cortex (Cheng, Liu, Shi, & Yan, 2017). Another category represented enriched pathways is chromatin and gene regulation which is characterized by a significant number (>100) of differentially expressed zinc finger proteins (ZNF). Variants in ZNF genes were previously linked to ADHD and other neurodevelopmental disorders (Al-Naama, Mackeh, & Kino, 2020).

One of the most notable findings of this study is the evidence of delayed neurite outgrowth in ADHD dopaminergic neurons. Differences in neurite outgrowth began to appear during the post-mitotic phase of differentiation (p = 0.008 at day 14), and by day 42 the neurites in the ADHD neurons had grown to reach full capacity, filling the culture well, and catching up to the comparison neurons. This delay in neuronal maturation in the ADHD**–**derived neurons support the theory of brain maturation delay in ADHD (Hoogman et al., 2017). This hypothesis suggests that the age at which peak cortical thickness is reached in the cerebrum is delayed in children with ADHD compared to neurotypical children (Shaw et al., 2007). Our RNA-seq data further support this observation. Gene set enrichment analysis revealed significant enrichment for biological functions related to neuronal development and differentiation. Notably, top significant DEGs such as *EPHA5* (logFC = –3.06, p = 1.03 x 10⁻⁹) and *PCDHA2* (logFC = – 6.02, p = 1.06 x 10⁻⁹) directly influence neurite outgrowth (Cooper et al., 2009; Shao et al., 2019). Additionally, ADHD-DEGs that overlap with genes implicated in ADHD-GWAS are overrepresented in neurodevelopmental and axonogenesis processes relevant to neurite outgrowth, indicating how transcriptomics can be effectively used to better unravel GWAS findings, supporting the observed neurite outgrowth delay in ADHD neurons.

Our transcriptomic findings, which demonstrated strong relevance to the aetiology of ADHD, underscore the importance of using a highly homogeneous and relevant cell type— dopaminergic neurons in our case—for comparative transcriptomic analyses. In a study using ADHD post-mortem brain tissue, while significant gene sets related to neurotransmitter activity (particularly glutamate) were identified, the heterogeneous nature of bulk brain tissue likely introduced high variability in RNA-seq results (Sudre et al., 2023). This could explain the relatively low number of significant DEGs identified in that study. Moreover, due to the limited availability of brain tissue, it is challenging to replicate such studies to confirm findings. Similarly, studies using blood samples—an irrelevant tissue for ADHD—have shown inconsistent results, with over 60% of DEGs identified in case-control comparisons failing to replicate in follow-up studies (Mortimer et al., 2020). Therefore, using a more homogeneous and relevant cell type can reduce variability, allowing consistent associations to be observed.

Despite the promising insights gained from this study, the examined sample size (one ADHD *vs* one comparison) may limit the generalizability of our findings (S. Zhao, Li, Guo, Sheng, & Shyr, 2018). A larger cohort that can account for inter**–** and intra**–**subject variability is critical to gain more definitive findings and avoid false positive/negative results (Rehbach, Fernando, & Brennand, 2020). In conclusion, this study represents a successful proof of principle that iPSCs can be useful in understanding the biological architecture of ADHD and other psychiatric conditions at a cellular and molecular level. Studying ADHD in the dopaminergic cell**–**type specific context using stem cell models and integrating it with transcriptomics offers a promising platform for future research and therapeutic development, paving the way for more effective and personalized avenues to managing ADHD. This approach can highlight key genetic and neurodevelopmental factors that contribute to the aetiology of this condition.

## References

1. Akutagava-Martins, G. C., Salatino-Oliveira, A., Genro, J. P., Contini, V., Polanczyk, G., Zeni, C., . . . Hutz, M.H. (2014). Glutamatergic copy number variants and their role in attention-deficit/hyperactivity disorder. Am J Med Genet B Neuropsychiatr Genet, 165B(6), 502–509. doi:10.1002/ajmg.b.32253

2. Al-Naama, N., Mackeh, R., & Kino, T. (2020). C(2)H(2)-Type Zinc Finger Proteins in Brain Development, Neurodevelopmental, and Other Neuropsychiatric Disorders: Systematic Literature-Based Analysis. Front Neurol, 11, 32. doi:10.3389/fneur.2020.00032

3. Allen-Brady, K., Miller, J., Matsunami, N., Stevens, J., Block, H., Farley, M., Coon, H. (2009). A high-density SNP genome-wide linkage scan in a large autism extended pedigree. Mol Psychiatry, 14(6), 590–600. doi:10.1038/mp.2008.14

4. Als, T. D., Kurki, M. I., Grove, J., Voloudakis, G., Therrien, K., Tasanko, E., Borglum, A. D. (2023). Depression pathophysiology, risk prediction of recurrence and comorbid psychiatric disorders using genome-wide analyses. Nat Med, 29(7), 1832–1844. doi:10.1038/s41591-023-02352-1

5. Arikkath, J., & Reichardt, L. F. (2008). Cadherins and catenins at synapses: roles in synaptogenesis and synaptic plasticity. Trends Neurosci, 31(9), 487–494. doi:10.1016/j.tins.2008.07.001

6. Ashburner, M., Ball, C. A., Blake, J. A., Botstein, D., Butler, H., Cherry, J. M., Sherlock, G. (2000). Gene ontology: tool for the unification of biology. The Gene Ontology Consortium. Nat Genet, 25(1), 25–29. doi:10.1038/75556

7. Barkley, R. A., & Fischer, M. (2019). Hyperactive Child Syndrome and Estimated Life Expectancy at Young Adult Follow-Up: The Role of ADHD Persistence and Other Potential Predictors. J Atten Disord, 23(9), 907–923. doi:10.1177/1087054718816164

8. Benjamini, Y., Drai, D., Elmer, G., Kafkafi, N., & Golani, I. (2001). Controlling the false discovery rate in behavior genetics research. Behav Brain Res, 125(1-2), 279–284. doi:10.1016/s0166-4328(01)00297-2

9. Berg, S., Kutra, D., Kroeger, T., Straehle, C. N., Kausler, B. X., Haubold, C., Kreshuk, A. (2019). ilastik: interactive machine learning for (bio)image analysis. Nat Methods, 16(12), 1226–1232. doi:10.1038/s41592-019-0582-9

10. Blauwendraat, C., Nalls, M. A., & Singleton, A. B. (2020). The genetic architecture of Parkinson’s disease. Lancet Neurol, 19(2), 170–178. doi:10.1016/S1474-4422(19)30287-X

11. Caiazzo, M., Dell’Anno, M. T., Dvoretskova, E., Lazarevic, D., Taverna, S., Leo, D., Broccoli, V. (2011). Direct generation of functional dopaminergic neurons from mouse and human fibroblasts. Nature, 476(7359), 224–227. doi:10.1038/nature10284

12. Cano-Gamez, E., & Trynka, G. (2020). From GWAS to Function: Using Functional Genomics to Identify the Mechanisms Underlying Complex Diseases. Front Genet, 11, 424. doi:10.3389/fgene.2020.00424

13. Cao, M., Park, D., Wu, Y., & De Camilli, P. (2020). Absence of Sac2/INPP5F enhances the phenotype of a Parkinson’s disease mutation of synaptojanin 1. Proc Natl Acad Sci U S A, 117(22), 12428–12434. doi:10.1073/pnas.2004335117

14. Cheng, J., Liu, A., Shi, M. Y., & Yan, Z. (2017). Disrupted Glutamatergic Transmission in Prefrontal Cortex Contributes to Behavioral Abnormality in an Animal Model of ADHD. Neuropsychopharmacology, 42(10), 2096–2104. doi:10.1038/npp.2017.30

15. Cherkasova, M. V., Roy, A., Molina, B. S. G., Scott, G., Weiss, G., Barkley, R. A., . . . Hechtman, L. (2022). Review: Adult Outcome as Seen Through Controlled Prospective Follow-up Studies of Children With Attention-Deficit/Hyperactivity Disorder Followed Into Adulthood. J Am Acad Child Adolesc Psychiatry, 61(3), 378–391. doi:10.1016/j.jaac.2021.05.019

16. Chou, M. Y., Hu, M. C., Chen, P. Y., Hsu, C. L., Lin, T. Y., Tan, M. J., . . . Huang, H. S. (2022). RTL1/PEG11 imprinted in human and mouse brain mediates anxiety-like and social behaviors and regulates neuronal excitability in the locus coeruleus. Hum Mol Genet, 31(18), 3161–3180. doi:10.1093/hmg/ddac110

17. Cooper, M. A., Kobayashi, K., & Zhou, R. (2009). Ephrin-A5 regulates the formation of the ascending midbrain dopaminergic pathways. Dev Neurobiol, 69(1), 36–46. doi:10.1002/dneu.20685

18. Cross-Disorder Group of the Psychiatric Genomics, C. (2013). Identification of risk loci with shared effects on five major psychiatric disorders: a genome-wide analysis. Lancet, 381(9875), 1371–1379. doi:10.1016/S0140-6736(12)62129-1

19. Dark, C., Homman-Ludiye, J., & Bryson-Richardson, R. J. (2018). The role of ADHD associated genes in neurodevelopment. Dev Biol, 438(2), 69–83. doi:10.1016/j.ydbio.2018.03.023

20. de Leeuw, C. A., Mooij, J. M., Heskes, T., & Posthuma, D. (2015). MAGMA: generalized gene-set analysis of GWAS data. PLoS Comput Biol, 11(4), e1004219. doi:10.1371/journal.pcbi.1004219

21. Demontis, D., Walters, G. B., Athanasiadis, G., Walters, R., Therrien, K., Nielsen, T. T., . . . Borglum, A. D. (2023). Genome-wide analyses of ADHD identify 27 risk loci, refine the genetic architecture and implicate several cognitive domains. Nat Genet, 55(2), 198–208. doi:10.1038/s41588-022-01285-8

22. Dobin, A., Davis, C. A., Schlesinger, F., Drenkow, J., Zaleski, C., Jha, S., . . . Gingeras, T. R. (2013). STAR: ultrafast universal RNA-seq aligner. Bioinformatics, 29(1), 15–21. doi:10.1093/bioinformatics/bts635

23. Elia, J., Glessner, J. T., Wang, K., Takahashi, N., Shtir, C. J., Hadley, D., . . . Hakonarson, H. (2011). Genome-wide copy number variation study associates metabotropic glutamate receptor gene networks with attention deficit hyperactivity disorder. Nat Genet, 44(1), 78–84. doi:10.1038/ng.1013

24. Faraone, S. V., Bellgrove, M. A., Brikell, I., Cortese, S., Hartman, C. A., Hollis, C., . . . Buitelaar, J. K. (2024). Attention-deficit/hyperactivity disorder. Nat Rev Dis Primers, 10(1), 11. doi:10.1038/s41572-024-00495-0

25. Faraone, S. V., & Larsson, H. (2019). Genetics of attention deficit hyperactivity disorder. Mol Psychiatry, 24(4), 562–575. doi:10.1038/s41380-018-0070-0

26. Fayyad, J., De Graaf, R., Kessler, R., Alonso, J., Angermeyer, M., Demyttenaere, K., . . . Jin, R. (2007). Cross-national prevalence and correlates of adult attention-deficit hyperactivity disorder. Br J Psychiatry, 190, 402–409. doi:10.1192/bjp.bp.106.034389

27. Felix Krueger, F. J., Phil Ewels, Ebrahim Afyounian, & Benjamin Schuster-Boeckler . . (2021). FelixKrueger/TrimGalore: v0.6.7 - DOI via Zenodo (0.6.7). Zenodo. 10.5281/zenodo.5127899

28. Ferranti, A. S., Luessen, D. J., & Niswender, C. M. (2024). Novel pharmacological targets for GABAergic dysfunction in ADHD. Neuropharmacology, 249, 109897. doi:10.1016/j.neuropharm.2024.109897

29. Fulton, S., Thibault, D., Mendez, J. A., Lahaie, N., Tirotta, E., Borrelli, E., . . . Trudeau, L. E. (2011). Contribution of Kv1.2 voltage-gated potassium channel to D2 autoreceptor regulation of axonal dopamine overflow. J Biol Chem, 286(11), 9360–9372. doi:10.1074/jbc.M110.153262

30. Garcia-Ortega, L. F., & Martinez, O. (2015). How Many Genes Are Expressed in a Transcriptome? Estimation and Results for RNA-Seq. PLoS One, 10(6), e0130262. doi:10.1371/journal.pone.0130262

31. Gillis, J., Mistry, M., & Pavlidis, P. (2010). Gene function analysis in complex data sets using ErmineJ. Nat Protoc, 5(6), 1148–1159. doi:10.1038/nprot.2010.78

32. Giros, B., Jaber, M., Jones, S. R., Wightman, R. M., & Caron, M. G. (1996). Hyperlocomotion and indifference to cocaine and amphetamine in mice lacking the dopamine transporter. Nature, 379(6566), 606–612. doi:10.1038/379606a0

33. Grove, J., Ripke, S., Als, T. D., Mattheisen, M., Walters, R. K., Won, H., . . . Borglum, A. D. (2019). Identification of common genetic risk variants for autism spectrum disorder. Nat Genet, 51(3), 431–444. doi:10.1038/s41588-019-0344-8

34. Hamada, N., Ito, H., Nishijo, T., Iwamoto, I., Morishita, R., Tabata, H., . . . Nagata, K. (2016). Essential role of the nuclear isoform of RBFOX1, a candidate gene for autism spectrum disorders, in the brain development. Sci Rep, 6, 30805. doi:10.1038/srep30805

35. Handunnetthi, L., Saatci, D., Hamley, J. C., & Knight, J. C. (2021). Maternal immune activation downregulates schizophrenia genes in the foetal mouse brain. Brain Commun, 3(4), fcab275. doi:10.1093/braincomms/fcab275

36. Harshil Patel, P. E., Alexander Peltzer, Olga Botvinnik, Gregor Sturm, Denis Moreno, Pranathi Vemuri, Silviamorins, Lorena Pantano, Mahesh Binzer-Panchal, nf-core bot, Gavin Kelly, Maxime U. Garcia, Friederike Hanssen, Matthias Zepper, James A. Fellows Yates, Chris Cheshire, rfenouil, Jose Espinosa-Carrasco, … George Hall. (2023). nf-core/rnaseq: nf-core/rnaseq v3.10.1 - Plastered Rhodium Rudolph (3.10.1). Zenodo. 10.5281/zenodo.7505987

37. Hawi, Z., Cummins, T. D., Tong, J., Johnson, B., Lau, R., Samarrai, W., & Bellgrove, M. A. (2015). The molecular genetic architecture of attention deficit hyperactivity disorder. Mol Psychiatry, 20(3), 289–297. doi:10.1038/mp.2014.183

38. Hawi, Z., Tong, J., Dark, C., Yates, H., Johnson, B., & Bellgrove, M. A. (2018). The role of cadherin genes in five major psychiatric disorders: A literature update. Am J Med Genet B Neuropsychiatr Genet, 177(2), 168–180. doi:10.1002/ajmg.b.32592

39. Hindley, G., Bahrami, S., Steen, N. E., O’Connell, K. S., Frei, O., Shadrin, A., . . . Andreassen, O. A. (2021). Characterising the shared genetic determinants of bipolar disorder, schizophrenia and risk-taking. Transl Psychiatry, 11(1), 466. doi:10.1038/s41398-021-01576-4

40. Hoogman, M., Bralten, J., Hibar, D. P., Mennes, M., Zwiers, M. P., Schweren, L. S. J., . . . Franke, B. (2017). Subcortical brain volume differences in participants with attention deficit hyperactivity disorder in children and adults: a cross-sectional mega-analysis. Lancet Psychiatry, 4(4), 310–319. doi:10.1016/S2215-0366(17)30049-4

41. Iida, I., Konno, K., Natsume, R., Abe, M., Watanabe, M., Sakimura, K., & Terunuma, M. (2024). Behavioral analysis of kainate receptor KO mice and the role of GluK3 subunit in anxiety. Sci Rep, 14(1), 4521. doi:10.1038/s41598-024-55063-z

42. Jerng, H. H., Lauver, A. D., & Pfaffinger, P. J. (2007). DPP10 splice variants are localized in distinct neuronal populations and act to differentially regulate the inactivation properties of Kv4-based ion channels. Mol Cell Neurosci, 35(4), 604–624. doi:10.1016/j.mcn.2007.03.008

43. Jones, R. M., Cadby, G., Melton, P. E., Abraham, L. J., Whitehouse, A. J., & Moses, E. K. (2013). Genome-wide association study of autistic-like traits in a general population study of young adults. Front Hum Neurosci, 7, 658. doi:10.3389/fnhum.2013.00658

44. Kampmann, M. (2020). CRISPR-based functional genomics for neurological disease. Nat Rev Neurol, 16(9), 465–480. doi:10.1038/s41582-020-0373-z

45. Kaulin, Y. A., De Santiago-Castillo, J. A., Rocha, C. A., Nadal, M. S., Rudy, B., & Covarrubias, M. (2009). The dipeptidyl-peptidase-like protein DPP6 determines the unitary conductance of neuronal Kv4.2 channels. J Neurosci, 29(10), 3242–3251. doi:10.1523/JNEUROSCI.4767-08.2009

46. Kitazawa, M., Sutani, A., Kaneko-Ishino, T., & Ishino, F. (2021). The role of eutherian-specific RTL1 in the nervous system and its implications for the Kagami-Ogata and Temple syndromes. Genes Cells, 26(3), 165–179. doi:10.1111/gtc.12830

47. Krause, K. H., Dresel, S. H., Krause, J., Kung, H. F., & Tatsch, K. (2000). Increased striatal dopamine transporter in adult patients with attention deficit hyperactivity disorder: effects of methylphenidate as measured by single photon emission computed tomography. Neurosci Lett, 285(2), 107–110. doi:10.1016/s0304-3940(00)01040-5

48. Lachman, H. M. (2008). Copy variations in schizophrenia and bipolar disorder. Cytogenet Genome Res, 123(1-4), 27–35. doi:10.1159/000184689

49. Law, C. W., Chen, Y., Shi, W., & Smyth, G. K. (2014). voom: Precision weights unlock linear model analysis tools for RNA-seq read counts. Genome Biol, 15(2), R29. doi:10.1186/gb-2014-15-2-r29

50. Lee, J. R. (2015). Protein tyrosine phosphatase PTPRT as a regulator of synaptic formation and neuronal development. BMB Rep, 48(5), 249–255. doi:10.5483/bmbrep.2015.48.5.037

51. Levy, F., Hay, D. A., McStephen, M., Wood, C., & Waldman, I. (1997). Attention-deficit hyperactivity disorder: a category or a continuum? Genetic analysis of a large-scale twin study. J Am Acad Child Adolesc Psychiatry, 36(6), 737–744. doi:10.1097/00004583-199706000-00009

52. Liao, C., Laporte, A. D., Spiegelman, D., Akcimen, F., Joober, R., Dion, P. A., & Rouleau, G. A. (2019). Transcriptome-wide association study of attention deficit hyperactivity disorder identifies associated genes and phenotypes. Nat Commun, 10(1), 4450. doi:10.1038/s41467-019-12450-9

53. Lim, S. H., Shin, S., Kim, M. H., Kim, E. C., Lee, D. Y., Moon, J., . . . Lee, J. R. (2020). Depression-like behaviors induced by defective PTPRT activity through dysregulated synaptic functions and neurogenesis. J Cell Sci, 133(20). doi:10.1242/jcs.243972

54. Longman, C., Brockington, M., Torelli, S., Jimenez-Mallebrera, C., Kennedy, C., Khalil, N., . . . Muntoni, F. (2003). Mutations in the human LARGE gene cause MDC1D, a novel form of congenital muscular dystrophy with severe mental retardation and abnormal glycosylation of alpha-dystroglycan. Hum Mol Genet, 12(21), 2853–2861. doi:10.1093/hmg/ddg307

55. Maffie, J., & Rudy, B. (2008). Weighing the evidence for a ternary protein complex mediating A-type K+ currents in neurons. J Physiol, 586(23), 5609–5623. doi:10.1113/jphysiol.2008.161620

56. Malloy, C., Ahern, M., Lin, L., & Hoffman, D. A. (2022). Neuronal Roles of the Multifunctional Protein Dipeptidyl Peptidase-like 6 (DPP6). Int J Mol Sci, 23(16). doi:10.3390/ijms23169184

57. Marshall, C. R., Noor, A., Vincent, J. B., Lionel, A. C., Feuk, L., Skaug, J., . . . Scherer, S.W. (2008). Structural variation of chromosomes in autism spectrum disorder. Am J Hum Genet, 82(2), 477–488. doi:10.1016/j.ajhg.2007.12.009

58. Matos, M. R., Ho, S. M., Schrode, N., & Brennand, K. J. (2020). Integration of CRISPR-engineering and hiPSC-based models of psychiatric genomics. Mol Cell Neurosci, 107, 103532. doi:10.1016/j.mcn.2020.103532

59. Mazzocchi, G., Andreis, P. G., De Caro, R., Aragona, F., Gottardo, L., & Nussdorfer, G. G. (1999). Cerebellin enhances in vitro secretory activity of human adrenal gland. J Clin Endocrinol Metab, 84(2), 632–635. doi:10.1210/jcem.84.2.5462

60. McCaffrey, T. A., St Laurent, G., 3rd, Shtokalo, D., Antonets, D., Vyatkin, Y., Jones, D., . . . Nigg, J. T. (2020). Biomarker discovery in attention deficit hyperactivity disorder: RNA sequencing of whole blood in discordant twin and case-controlled cohorts. BMC Med Genomics, 13(1), 160. doi:10.1186/s12920-020-00808-8

61. Mortimer, N., Sanchez-Mora, C., Rovira, P., Vilar-Ribo, L., Richarte, V., Corrales, M., . . . Ribases, M. (2020). Transcriptome profiling in adult attention-deficit hyperactivity disorder. Eur Neuropsychopharmacol, 41, 160–166. doi:10.1016/j.euroneuro.2020.11.005

62. Mullins, N., Forstner, A. J., O’Connell, K. S., Coombes, B., Coleman, J. R. I., Qiao, Z., . . . Andreassen, O.A. (2021). Genome-wide association study of more than 40,000 bipolar disorder cases provides new insights into the underlying biology. Nat Genet, 53(6), 817–829. doi:10.1038/s41588-021-00857-4

63. Muzes, G., & Sipos, F. (2016). Heterogeneity of Stem Cells: A Brief Overview. Methods Mol Biol, 1516, 1–12. doi:10.1007/7651_2016_345

64. Nakatsu, F., Messa, M., Nandez, R., Czapla, H., Zou, Y., Strittmatter, S. M., & De Camilli, P. (2015). Sac2/INPP5F is an inositol 4-phosphatase that functions in the endocytic pathway. J Cell Biol, 209(1), 85–95. doi:10.1083/jcb.201409064

65. Nalls, M. A., Pankratz, N., Lill, C. M., Do, C. B., Hernandez, D. G., Saad, M., . . . Singleton, A.B. (2014). Large-scale meta-analysis of genome-wide association data identifies six new risk loci for Parkinson’s disease. Nat Genet, 46(9), 989–993. doi:10.1038/ng.3043

66. Namipashaki, A., Pugsley, K., Liu, X., Abrehart, K., Lim, S. M., Sun, G., . . . Hawi, Z. (2023). Integration of xeno-free single-cell cloning in CRISPR-mediated DNA editing of human iPSCs improves homogeneity and methodological efficiency of cellular disease modeling. Stem Cell Reports, 18(12), 2515–2527. doi:10.1016/j.stemcr.2023.10.013

67. Nolbrant, S., Heuer, A., Parmar, M., & Kirkeby, A. (2017). Generation of high-purity human ventral midbrain dopaminergic progenitors for in vitro maturation and intracerebral transplantation. Nat Protoc, 12(9), 1962–1979. doi:10.1038/nprot.2017.078

68. Noor, A., Lionel, A. C., Cohen-Woods, S., Moghimi, N., Rucker, J., Fennell, A., . . . Vincent, J.B. (2014). Copy number variant study of bipolar disorder in Canadian and UK populations implicates synaptic genes. Am J Med Genet B Neuropsychiatr Genet, *165B*(4), 303-313. doi:10.1002/ajmg.b.32232

69. O’Leary, A., Fernandez-Castillo, N., Gan, G., Yang, Y., Yotova, A. Y., Kranz, T. M., . . . Reif, A. (2022). Behavioural and functional evidence revealing the role of RBFOX1 variation in multiple psychiatric disorders and traits. Mol Psychiatry, 27(11), 4464–4473. doi:10.1038/s41380-022-01722-4

70. Olgiati, S., De Rosa, A., Quadri, M., Criscuolo, C., Breedveld, G. J., Picillo, M., . . . Bonifati, V. (2014). PARK20 caused by SYNJ1 homozygous Arg258Gln mutation in a new Italian family. Neurogenetics, 15(3), 183–188. doi:10.1007/s10048-014-0406-0

71. Patro, R., Duggal, G., Love, M. I., Irizarry, R. A., & Kingsford, C. (2017). Salmon provides fast and bias-aware quantification of transcript expression. Nat Methods, 14(4), 417–419. doi:10.1038/nmeth.4197

72. Powell, S. K., O’Shea, C., Townsley, K., Prytkova, I., Dobrindt, K., Elahi, R., . . . Brennand, K.J. (2023). Induction of dopaminergic neurons for neuronal subtype-specific modeling of psychiatric disease risk. Mol Psychiatry, 28(5), 1970–1982. doi:10.1038/s41380-021-01273-0

73. Powell, D. (2019). drpowell/degust 4.1.1 (4.1.1). Zenodo. 10.5281/zenodo.3501067

74. Raghavan, N. S., Dumitrescu, L., Mormino, E., Mahoney, E. R., Lee, A. J., Gao, Y., . . . Alzheimer’s Disease Neuroimaging, I. (2020). Association Between Common Variants in RBFOX1, an RNA-Binding Protein, and Brain Amyloidosis in Early and Preclinical Alzheimer Disease. JAMA Neurol, 77(10), 1288–1298. doi:10.1001/jamaneurol.2020.1760

75. Rehbach, K., Fernando, M. B., & Brennand, K. J. (2020). Integrating CRISPR Engineering and hiPSC-Derived 2D Disease Modeling Systems. J Neurosci, 40(6), 1176–1185. doi:10.1523/JNEUROSCI.0518-19.2019

76. Schindelin, J., Arganda-Carreras, I., Frise, E., Kaynig, V., Longair, M., Pietzsch, T., . . . Cardona, A. (2012). Fiji: an open-source platform for biological-image analysis. Nat Methods, 9(7), 676–682. doi:10.1038/nmeth.2019

77. Selten, M., van Bokhoven, H., & Nadif Kasri, N. (2018). Inhibitory control of the excitatory/inhibitory balance in psychiatric disorders. F1000Res, 7, 23. doi:10.12688/f1000research.12155.1

78. Shao, Z., Noh, H., Bin Kim, W., Ni, P., Nguyen, C., Cote, S. E., . . . Chung, S. (2019). Dysregulated protocadherin-pathway activity as an intrinsic defect in induced pluripotent stem cell-derived cortical interneurons from subjects with schizophrenia. Nat Neurosci, 22(2), 229–242. doi:10.1038/s41593-018-0313-z

79. Shaw, P., Eckstrand, K., Sharp, W., Blumenthal, J., Lerch, J. P., Greenstein, D., . . . Rapoport, J.L. (2007). Attention-deficit/hyperactivity disorder is characterized by a delay in cortical maturation. Proc Natl Acad Sci U S A, 104(49), 19649–19654. doi:10.1073/pnas.0707741104

80. Soneson, C., Love, M. I., & Robinson, M. D. (2015). Differential analyses for RNA-seq: transcript-level estimates improve gene-level inferences. F1000Res, 4, 1521. doi:10.12688/f1000research.7563.2

81. Sudre, G., Gildea, D. E., Shastri, G. G., Sharp, W., Jung, B., Xu, Q., . . . Shaw, P. (2023). Mapping the cortico-striatal transcriptome in attention deficit hyperactivity disorder. Mol Psychiatry, 28(2), 792–800. doi:10.1038/s41380-022-01844-9

82. Szklarczyk, D., Kirsch, R., Koutrouli, M., Nastou, K., Mehryary, F., Hachilif, R., . . . von Mering, C. (2023). The STRING database in 2023: protein-protein association networks and functional enrichment analyses for any sequenced genome of interest. Nucleic Acids Res, 51(D1), D638–D646. doi:10.1093/nar/gkac1000

83. Tong, J., Lee, K. M., Liu, X., Nefzger, C. M., Vijayakumar, P., Hawi, Z., . . . Bellgrove, M.A. (2019). Generation of four iPSC lines from peripheral blood mononuclear cells (PBMCs) of an attention deficit hyperactivity disorder (ADHD) individual and a healthy sibling in an Australia-Caucasian family. Stem Cell Res, 34, 101353. doi:10.1016/j.scr.2018.11.014

84. Trubetskoy, V., Pardinas, A. F., Qi, T., Panagiotaropoulou, G., Awasthi, S., Bigdeli, T. B., . . . Schizophrenia Working Group of the Psychiatric Genomics, C. (2022). Mapping genomic loci implicates genes and synaptic biology in schizophrenia. Nature, 604(7906), 502–508. doi:10.1038/s41586-022-04434-5

85. Zhao, S., Li, C. I., Guo, Y., Sheng, Q., & Shyr, Y. (2018). RnaSeqSampleSize: real data based sample size estimation for RNA sequencing. BMC Bioinformatics, 19(1), 191. doi:10.1186/s12859-018-2191-5

86. Zhao, W. W. (2013). Intragenic deletion of RBFOX1 associated with neurodevelopmental/neuropsychiatric disorders and possibly other clinical presentations. Mol Cytogenet, 6(1), 26. doi:10.1186/1755-8166-6-26

87. Zoubarev, A., Hamer, K. M., Keshav, K. D., McCarthy, E. L., Santos, J. R., Van Rossum, T., . . . Pavlidis, P. (2012). Gemma: a resource for the reuse, sharing and meta-analysis of expression profiling data. Bioinformatics, 28(17), 2272–2273. doi:10.1093/bioinformatics/bts430

